# Structural insights into abscisic acid exporter AtABCG25

**DOI:** 10.1101/2023.09.22.559085

**Authors:** Jian Xin, Yeling Zhou, Yichun Qiu, He Geng, Yuzhu Wang, Yi Song, Jiansheng Liang, Kaige Yan

**Affiliations:** Department of Chemical Biology, School of Life Sciences, Southern University of Science and Technology, Shenzhen 518055, China; Institute of Plant and Food Science, Department of Biology, School of Life Sciences, Southern University of Science and Technology, Shenzhen 518055, China; Max Planck Institute of Molecular Plant Physiology, Potsdam-Golm, Germany; Institute for Biological Electron Microscopy, Southern University of Science and Technology, Shenzhen 518055, China

**Author notes:** Correspondence: J. L., Y. S. and K.Y. Co-first authors (Jian Xin, Yeling Zhou, Yichun Qiu).

**Keywords:** AtABCG25, abscisic acid, ABC transporters, cryo-EM, structure

## Abstract

Cellular hormone homeostasis is essential for the precise spatial and temporal signaling responses and plant fitness. Abscisic acid (ABA) plays pivotal roles in orchestrating various developmental and stress responses and confers fitness benefits over ecological and evolutionary timescales in terrestrial plants. Cellular ABA levels is regulated by complex processes including biosynthesis, catabolism, and transport. AtABCG25 is the first identified ABA exporter through genetic screen which affects diverse ABA responses. Resolving the structure basis of ABCG25 in ABA exporting is critical for further manipulating ABA homeostasis and plant fitness. We utilized cryo-electron microscopy to elucidate the structural dynamics of AtABCG25, and successfully characterized different states including apo AtABCG25, ABA-bound AtABCG25 and ATP-bound AtABCG25(E232Q). Notably, AtABCG25 forms a homodimer, featuring a deep, slit-like cavity in the transmembrane domain. The critical residues in the cavity where ABA binds are precisely characterized. Moreover, ATP binding triggers the closure of nucleotide-binding domains and conformational transitions in the transmembrane domains. Collectively, these findings provide valuable insights into the intricate substrate recognition and transport mechanisms of ABA exporter ABCG25, paving the way towards genetical manipulating of ABA homeostasis and plant fitness.

## Introduction

Plant hormone signaling plays central roles in integrating environmental cues and internal developmental and stress responses (Spartz and Gray, 2008). ABA (abscisic acid) is a versatile hormone broadly regulates plant responses to abiotic and biotic stresses (Hu et al., 2022; Nakashima and Yamaguchi-Shinozaki, 2013; Zhao et al., 2016), as well as developmental processes like seed germination, leaf senescence and fruit maturation (Ali et al., 2022; Gao et al., 2016; Sun et al., 2017; Wang et al., 2013). ABA signaling perception pathway is largely conserved in land plants, which ABA binds to its receptor PYRABACTIN RESISTANCE 1-like (PYLs) to suppress PROTEIN PHOSPHATASE 2C (PP2C) activity, further leads to SNF1-RELATED PROTEIN KINASE2 (SnRK2) auto-activation to regulate downstream responses (Raghavendra et al., 2010). Interestingly, although ABA biosynthetic pathway is present in algae, ABA mediated regulation of PYL activity was acquired in the common ancestor of terrestrial plants (Sun et al., 2020). This further indicates the crucial roles of ABA signaling in adapting to terrestrial ecosystems in land plants. Several studies had resolved the structure of ABA receptor, which substantially furthered our understanding about the mechanism underlying ABA perception (Ali et al., 2022; Melcher et al., 2010). However, much less study focused on the structural basis of ABA transportation in plants.

As a stress responsive hormone, cellar ABA levels must be tightly regulated to maintain a proper response. Plants evolved both ABA exporters and importers to maintain ABA homeostasis. Most of those transporters belong to ATP-binding cassette (ABC) G family proteins, including the exporters like AtABCG25/31 and importers AtABCG17/18/30/40 (Zhang et al., 2023). ATP-binding cassette (ABC) transporters are a large family of transmembrane proteins involved in the transport of wide variety of substrates ranging from small ions to polypeptides or polysaccharides (Velamakanni et al., 2007). As integral membrane transporters, ABC proteins are generally featured with set of structural modules containing a hydrophobic transmembrane domain (TMD) and a cytosolic domain known as nucleotide-binding domain (NBD) (Jasinski et al., 2003). Depending on the number of core domain, ABC proteins are classified as half-size transporters (one TMD-NBD) and full-size transporters (two TMD-NBDs) (Tarr et al., 2009). Notably, half-size transporters are generally more prevailed than full-size transporters in many land plants and animals (Hwang et al., 2016). Besides, higher plants possess two-fold more ABC transporters than other eukaryotes and particularly enriched in the B and G subfamilies (Do et al., 2021; Hwang et al., 2016). Investigating the structures of ABC transporters would help reveal their essential roles in human disease and plant health.

As the first genetically identified ABA exporter, AtABCG25 is a critical transporter affecting diverse ABA responses (Kuromori et al., 2010). For instance, AtABCG25 collaborates with ABCG31 to export ABA from the endosperm to the embryo, thereby regulating seed germination (Kang et al., 2015). AtABCG25 plays a role in stomatal regulation, and overexpression of ABCG25 leads to an increase in leaf temperature and a reduction in water loss rate (Kuromori et al., 2016). For other ABA transporters, AtABCG40 was an ABA importer in *Arabidopsis*, regulating stomatal aperture and drought tolerance (Kang et al., 2010). AtABCG30 and AtABCG40 showed ABA influx activity, facilitating ABA into the embryo, where seed dormancy occurs (Kang et al., 2015). In contrast, AtABCG17 and AtABCG18 act redundantly to boost ABA import and ultimately impede stomatal closure (Zhang et al., 2021). Despite that increasing evidence documenting the crucial roles of ABA transporters in ABA responses, there is a scarcity of molecular insights into its substrate selectivity and transport mechanisms.

In this study, we determined high-resolution structures of Arabidopsis ABCG25 using cryo-EM, encompassing apo, ABA-bound, and ATP-bound states. Our comprehensive structural analyses not only elucidate the architectural details of ABCG25 but also provide insights into ABA recognition and subsequent conformational changes during ABA transport.

## Result

### Purification and ATPase activity of AtABCG25

To investigate the molecular mechanism of ABA transport by AtABCG25, we conducted recombinant expression of AtABCG25, with a 3× Flag tag fused to its N-terminus. The protein was expressed by using the *Spodoptera frugiperda 9* (Sf9) insect cells. 1% Lauryl Maltose Neopentyl Glycol (LMNG) and 0.2% cholesteryl hemisuccinate (CHS) were used to extract AtABCG25 from the cell membrane, and subsequently 0.02% glyco-diosgenin (GDN) were added to the wash and elution buffer in affinity purification and gel filtration. The protein exhibiting the best behavior was applied for cryo-EM single particle analysis (Supplemental Figure 1A). For apo state of AtABCG25, the CHS was removed in the extraction. We determined the ATPase activity in a buffer containing 0.02% GDN. AtABCG25, extracted with CHS, exhibited an ATPase activity with a *V*_max_ (maximal velocity) of 4474 nmolPi mg^-1^ h^-1^ and an apparent *K*_m_ (Michaelis constant) of 1.67 mM (Figure 1A). In the case of AtABCG25 extracted without CHS, the *V*_max_ was 3613 nmolPi mg^-1^ h^-1^ and the *K*_m_ was 2.96 mM. However, the Walker-B mutation E232Q did not exhibit ATPase activity. These results suggest that CHS may serve as a substrate for AtABCG25, thereby accelerating the enzymatic reaction.

**Figure 1.**
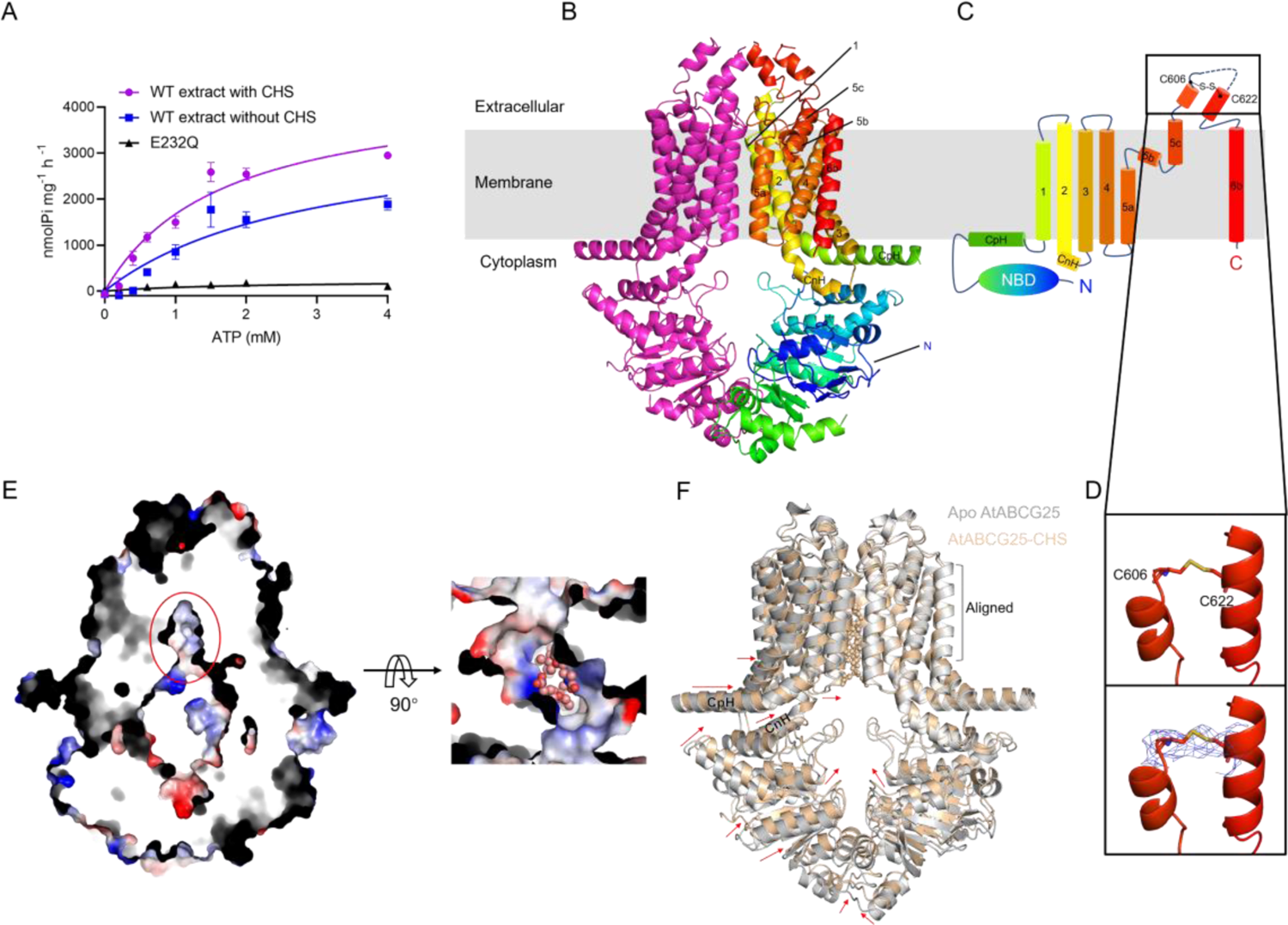
ATPase activity and overall architecture AtABCG25. (A) ATPase activities of both AtABCG25 wild type (WT) and the E232Q mutant. Data points represent the mean ± SD of three measurements. Data points were nonlinear-fitted using the Michaelis–Menten equation. (B) Overall structure of AtABCG25. One of the AtABCG25 monomers is depicted in a magenta color, while the other AtABCG25 monomer is represented with a color gradient spanning the rainbow spectrum, transitioning from blue to red. (C) The topological representation of the ABCG2 TMD follows the color scheme as depicted in Figure B. Cysteine residues involved in the formation of intramolecular disulfide bonds (C606 and C622) are specifically highlighted. (D) The details of intramolecular disulfide bonds (C606 and C622). (E) Sectional view depicting the surface electrostatic potential of CHS-bound AtABCG25 from the membrane’s side. Anionic and cationic charges are respectively depicted in red and blue. CHS is depicted in stick-and-ball representation. (F) Structural alignment between apo AtABCG25 and CHS-bound AtABCG25. Apo AtABCG25 is depicted in gray, while CHS-bound AtABCG25 is shown in wheat. CHS is depicted in stick-and-ball representation.

### Overall architecture of AtABCG25

The AtABCG25 cryo-EM map was reconstructed at an overall resolution of 3.05 Å (Fourier shell correlation (FSC) = 0.143 criterion) (Supplemental Figure 2A). As a half-size transporter, AtABCG25 functions in homodimer form. The folding of AtABCG25 resembles that of the human multidrug resistance protein hsABCG2. Compared to the ABCB subfamily, AtABCG25 exhibits the same short transmembrane helices and intracellular loops as the other ABCG subfamily proteins, resulting in a smaller distance between the NBD and the TMD (Supplemental Figure 4C). This makes AtABCG25 a more compact ABC transporter protein. The TMD consists of six transmembrane helices (TM1-TM6). There is a connecting helix (CnH) located just before TM1, nearly parallel to the membrane. A coupling helix (CpH) connects TM2 and TM3 (Figure 1B and 1C). The TMD interface of AtABCG25 is formed by the relative monomers of TM1, TM2, and TM5a. The structures of AtABCG25 and hsABCG2 are similar, with an RMSD of 2.86 Å between them (Supplemental Figure 4D). Despite the absence of bound nucleotides, the NBD of AtABCG25 remains in contact. Like human hsABCG2, intramolecular disulfide bonds (C606-C622 and C606’-C622’) are present in AtABCG25 (Figure 1D). The intramolecular disulfide bonds in hsABCG2 are important for structural stability and activity. However, the intermolecular disulfide bonds that are present in human ABCG2 are absent in AtABCG25. As the intermolecular disulfide bond in hsABCG2 was not required for cellular transport function (Kage et al., 2005), the lack of intermolecular disulfide bond in AtABCG25 seems to have little influence on its transport of ABA.

The overall architecture of AtABCG25 reveals an inward-opening conformation with a deep, slit-like cavity formed primarily by TM1, TM2, and TM5a from two AtABCG25 monomers (Figure 1E). The cavity is oriented towards the cytoplasm and extends approximately halfway through the membrane’s depth. Two large density features symmetrically located in the cavity were observed, whose sizes and shapes matched extremely well with the density of the cholesterol analogue CHS that had been added during purification. Since no other lipid molecules was added during the purification process, we interpreted the densities as two CHS molecules (Figure 1E and Supplemental Figure 3A). The shape of the densities guided us to place the CHS with their hydroxyl groups away from the cavity, in the opposite direction of the bound cholesterol in hsABCG2. The C2 symmetry was used during data processing, and processing with C1 symmetry gave similar results. Cholesterol is not a bona fide substrate of hABCG2 but can still bind to hABCG2. Similarly, the cholesterol analogue CHS is not a substrate of AtABCG25 but can still bind to AtABCG25. Human hABCG2 did not catalyze significant cholesterol transport *in vivo*, but its function has been shown to be regulated by cholesterol (Telbisz et al., 2014). It is possible that AtABCG25 function is also regulated by some lipid molecules.

In addition, we also obtained the apo state of AtABCG25, the CHS was removed in the extraction. The AtABCG25 without CHS cryo-EM map was reconstructed at an overall resolution of 2.87 Å (FSC = 0.143 criterion) (Supplemental Figure 2B). There is also unidentified density in the cavity of apo AtABCG25 (Supplemental Figure 3B). This density could potentially correspond to sterols, which might be extracted from Sf9 cells. Compared to CHS-bound AtABCG25, the apo AtABCG25 without CHS reveals a more open conformation. The combination of CHS makes the TMD of AtABCG25 close to each other, further driving the two NBD domains in tight contact. CpH and CnH also display significant proximity (Figure 1F).

### Substrate-specificity and auxiliary binding sites

For ABA bound cryo-EM sample preparation, 0.2 mM ABA was introduced into the AtABCG25 prior to cryo-EM analysis. The ABA-bound AtABCG25 map was refined to an overall resolution of 3.23 Å, and both the TMD as well as the substrate-binding cavity could be clearly resolved (Figure 2A and Supplemental Figure 2C and 3C). We observe the density features in the substrate-binding cavity, which is also formed by the transmembrane helices TM1, TM2, and TM5a of the opposing AtABCG25 monomers. The density within the cavity appears to accommodate only one ABA molecule. ABA can bind in either direction regardless of a 180° rotation, given that AtABCG25 exhibits C2 symmetry (Figure 2B and Supplemental Figure 3C). However, two ABA molecules cannot bind at the same time because their rings would spatially clash. The strongest density occurs at the axis of symmetry, where the core of the ABA flattened ring binds. No significant conformational changes were observed in the comparison between ABA-bound AtABCG25 and CHS-bound AtABCG25 (Figure 2C). This might be attributed by the fact that the CHS binding state corresponds to a considerably more compacted TMD in AtABCG25. Comparing to the apo AtABCG25, it is evident that a significant conformational change occurs in ABA-bound AtABCG25 (Figure 2D and Supplemental Figure 4B). The presence of ABA results in the convergence of the whole TMD of AtABCG25, subsequently bringing the NBD into proximity. Additionally, CpH and CnH also exhibits a notable level of proximity (Figure 2D).

**Figure 2.**
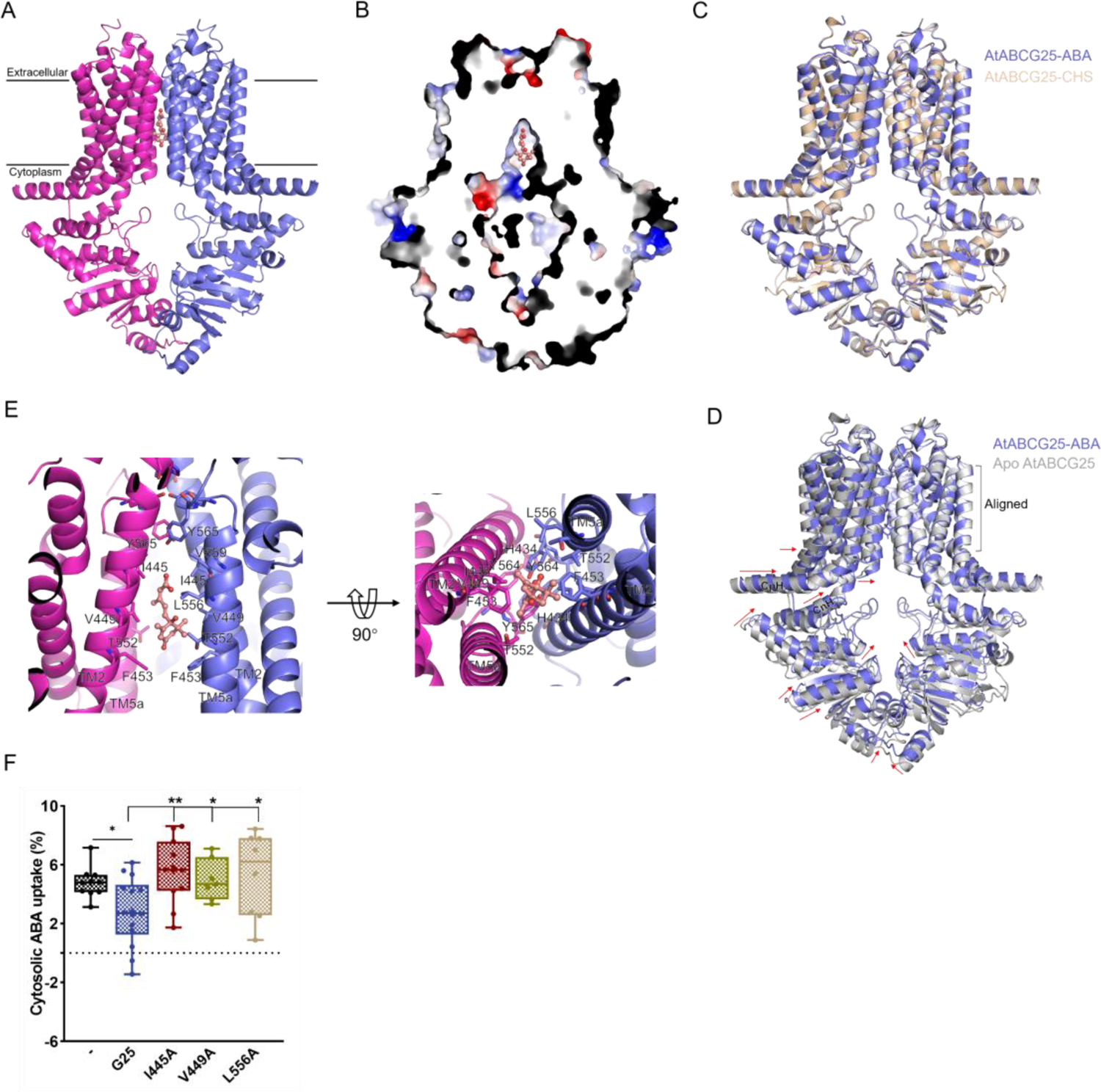
The structure of AtABCG25 bound to ABA. (A) Overall structure of AtABCG25 bound to ABA. ABA is represented in a stick-and-ball form. (B) Sectional view depicting the surface electrostatic potential of ABA-bound AtABCG25 from the membrane’s side. Anionic and cationic charges are respectively depicted in red and blue. ABA binds within the cavity. (C) Structural alignment between CHS-bound AtABCG25 and ABA-bound AtABCG25. CHS-bound AtABCG25 is depicted in wheat, while ABA-bound AtABCG25 is shown in slate. Residues not used are hidden. (D) Structural alignment between apo AtABCG25 and ABA-bound AtABCG25. Apo AtABCG25 is depicted in gray, while ABA-bound AtABCG25 is shown in slate. Residues not used are hidden. (E) The side and bottom view of the ABA-binding site. (F) Characterization of ABA transport activity for ABCG25 wild type and mutants in tobacco mesophyll protoplasts. Tobacco cells were separated into groups that were either not transfected (control group, -) or transfected with ABCG25 wild type, G25 mutants I445A, V449A, L556A. Shown are cellular ABA uptake characterized as relative cytosolic ABA level (ABA/mock treatment) and presented as box plots with original data points. Statistics were performed using one-way ANOVA with Dunnett’s multiple comparisons test (**\*\****P*<0.05, **\****P*<0.05 for control group versus WT and G25 mutants versus WT).

**Figure 3.**
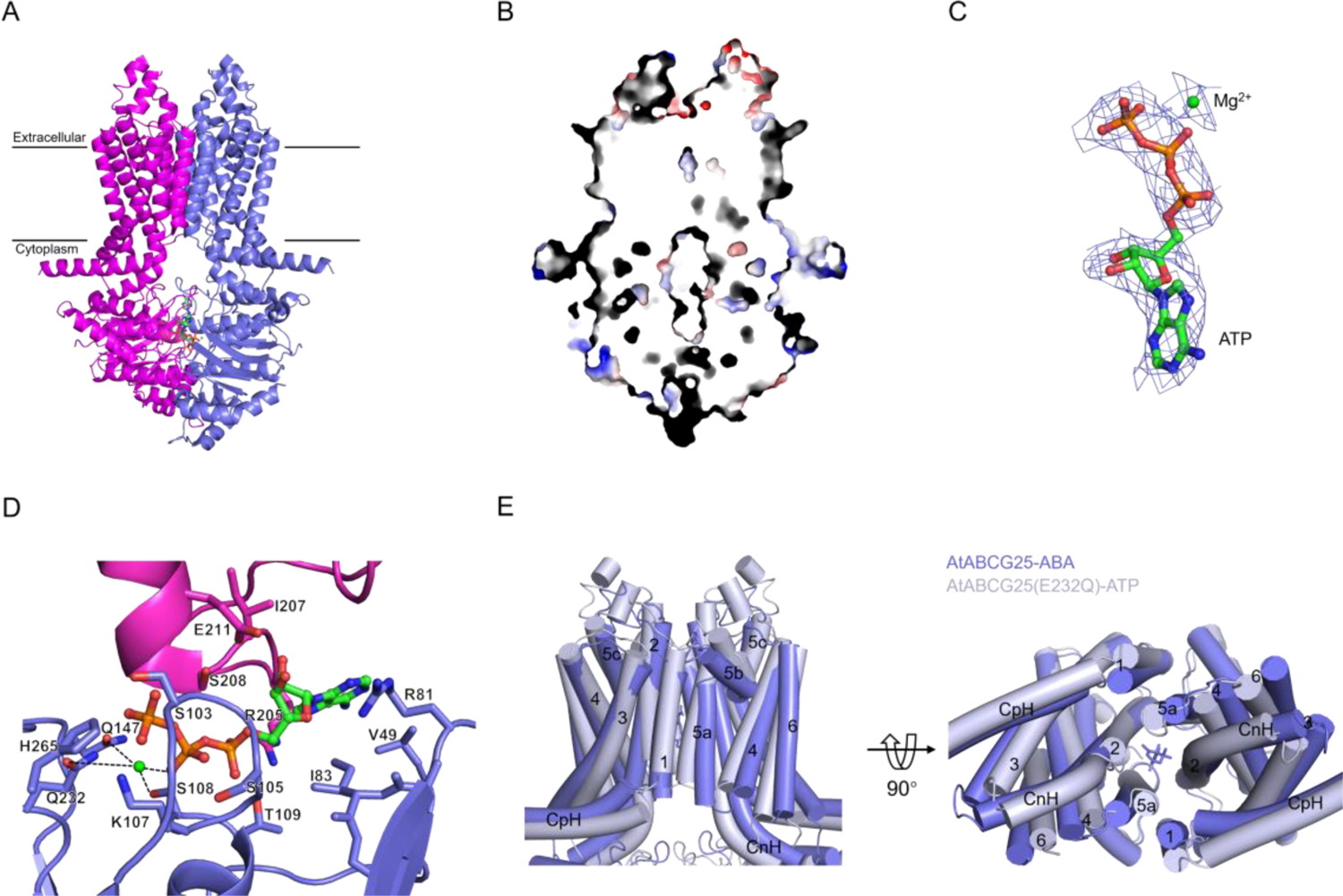
The structure of ATP-bound AtABCG25(E232Q). (A) Overall outward structure of AtABCG25(E232Q) bound to ATP and magnesium. ATP is represented in a stick-and-ball form. In the structure, nitrogen atoms of ATP molecules are depicted in blue, phosphorus atoms in orange, and magnesium ions in red. (B) Sectional view depicting the surface electrostatic potential of ABA-bound AtABCG25 from the membrane’s side. Anionic and cationic charges are respectively depicted in red and blue. ABA binds within the cavity. (C) Density map of ATP and Mg^2+^ in AtABCG25 outward. (D) The detailed structure of ATP binding site. One of the AtABCG25(E232Q) monomers is depicted in magenta, while the other AtABCG25(E232Q) monomer is represented with a slate color. (E) Structural alignment between ATP-bound AtABCG25(E232Q) and ABA-bound AtABCG25. ATP-bound AtABCG25(E232Q) is depicted in bluewhite, while ABA-bound AtABCG25 is shown in slate. Residues not used are hidden.

**Figure 4.**
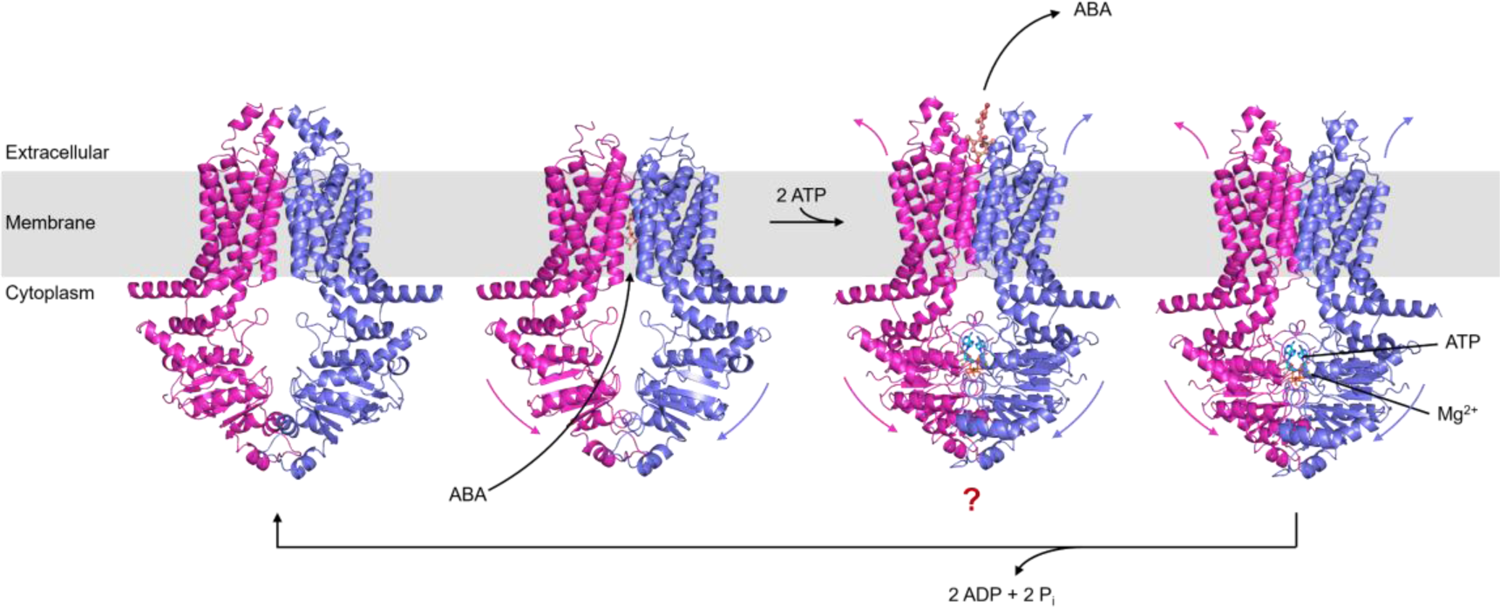
The ATP-driven cycle in ABCG25 for transporting ABA. AtABCG25 transports ABA through an ATP-driven process. In its apo state, the TMD of AtABCG25 forms a cavity with minimal interaction with the NBD. Upon ABA entry into the cytoplasmic cavity, it binds to AtABCG25, bringing the NBDs closer together. Subsequently, ATP binding further narrows the NBD distance. As the NBDs approach, the TMD’s cytoplasmic regions converge, while the extracellular part moves apart, facilitating ABA release. When NBD proximity allows ATP hydrolysis, it converts into ADP and Pi, which are then released, leading AtABCG25 back to its apo state.

ABA binds to AtABCG25 primarily through hydrophobic interactions. The carboxylate group of ABA is located at the bottom of the cavity of AtABCG25 homodimer, with the cyclohexene group backing away from the bottom of the cavity (Figure 2B and Supplemental Figure 3C). Moreover, ABA forms hydrophobic interactions primarily with hydrophobic amino acids possessing long side chains, comprising a combination of residues I445, V449, F453, T552, L556, and V559 on one TMD, along with I445, V449, F453, and T552 in the other TMD (Figure 2E). All these amino acids are located on TM2 and TM5a of the TMDs, with I445, V449, F453 on TM2 and T552, L556, V559 on TM5a. Y565, Y564, H434 are located at the bottom of the cavity and are in the closed state (Figure 2E). In addition, the residues Q431, F457and K545 are also involved in the cavity formation. To further determine the functions of the main residues forming ABA binding cavity, we generated several ABCG25 variants with each of the residues mutated to alanine (I445A, V449A, L556A) and tested their ABA transport activities using the ABA sensor-based tobacco cell assay (Zhou et al., 2021). As expected, treatment of tobacco cells with ABA resulted in an increase in the cytosolic ABA level, whereas co-transfected with ABCG25 led to a notable reduction in cellular ABA uptake (Figure 2F), validating the ABA efflux activity of ABCG25. Nonetheless, cells co-transfected with all the three variants exhibited cellular ABA uptake activities that were similar to those cells without transporter transfection (Figure 2F), indicating a loss of ABA efflux activity in the variant groups. These results suggest that all the three residues in ABCG25 are essential for its ABA transport activity.

### ATP and magnesium binding sites of AtABCG25(E232Q)

To further elucidate the ABA efflux mechanism of AtABCG25, obtaining an “outward” conformation of AtABCG25 is essential. The introduction of the E232Q mutation in AtABCG25 resulted in a nearly complete abolishment of ATP hydrolysis activity, allowing for the effective stabilization of AtABCG25 in the ATP-bound state. By adding ATP, Mg^2+^, and ABA to the AtABCG25(E232Q) protein for cryo-sampling, we obtained AtABCG25(E232Q) with an ATP-bound conformation in which the NBD dimer is tightly closed (Figure 3A-3B and Supplemental Figure 3D). The AtABCG25(E232Q) cryo-EM map was reconstructed at an overall resolution of 3.3 Å (Supplemental Figure 2D). The structure unveils an NBD dimer in a tightly closed conformation, with ATP binding occurring between the Walker A motif (GPSGSGKST) of one NBD and the signature motif (ISGGE) of the other NBD (Figure 3D). In addition to the ATP molecule being bound (Figure 3C-3D), Mg^2+^ is also coordinated with it. The coordination of the γ-phosphate of ATP involves three conserved side chains, namely Q232 (corresponding to the catalytic glutamate in wild type AtABCG25), H265 (functioning as the pivotal “switch” histidine), and Q147 (a constituent of the Q-loop) (Figure 3D). Q232 also coordinates the magnesium ion. On the ATP contact surface, one face of the ATP adenine ring is within Van der Waals distance of an NBD, with the residues I83 and V49, R81 and H265 forming salt bonds with ATP.

### Conformational dynamics associated with the export of ABA by AtABCG25

As a result of ATP binding, the NBD structural domains move closer to each other and come into closer contact. The NBD of each ABCG25 monomer undergoes an inward swing relative to the ATP unbound state. As a result of the movement of the NBDs, the cytoplasmic portions of the TMD are pushed toward each other, and the extracellular portion of the TMD produces a partial opening. The conformational change of the TMD is akin to a rigid-body movement (Figure 3E and Supplemental Figure 4A). The CpH and CnH of the AtABCG25 appear to be significantly closer to each other. On the extracellular sides of TM5a and TM2, there is a separation, creating a pathway for the export of ABA. Conversely, on the intracellular sides of TM5a and TM2, there is a closer approach, leading to the closure of the original cavities. In the ABA-bound AtABCG25 conformation, Y565, Y564, and H434 are in the closed state, with 3.5 Å between Y564 and Y564’, 2.7 Å between Y565 and Y565’, and 3.1 Å between H434 and H434’ (Supplemental Figure 5A). In the outward conformation, the closed state transitions to an open state, and the distance between them increases, with the distance between Y564 and Y564’ increasing to 4.4 Å, Y565 and Y565’ to 3.8 Å, and H434 and H434’ to 4.6 Å (Supplemental Figure 5B).

The transport of ABA by AtABCG25 is driven by ATP. In the apo state, the TMD of AtABCG25 forms a cavity with minimal contact with the NBD. When ABA enters the cavity of AtABCG25 from the cytoplasm, it binds to the cavity of AtABCG25. ABA binding brings the NBD of AtABCG25 closer together. Subsequently, ATP binds to the NBD of AtABCG25, and ATP binding further reduces the distance between the NBDs. As the NBDs approach, the cytoplasmic portion of the TMD moves closer to each other, while the extracellular portion of the TMD moves away, allowing ABA to be released outside the cell. When the NBDs reach a proximity conducive to ATP hydrolysis, ATP undergoes hydrolysis, resulting in the production and release of ADP and Pi. Subsequently, AtABCG25 reverts to the apo state.

## Discussion

Our structural analyses unveil the distinct conformations of AtABCG25 in its apo, ABA-bound, and ATP-bound states, providing insights into the molecular mechanisms underpinning ABA recognition and the essential conformational changes facilitating ABA transport. A notable observation in the apo-state structure of AtABCG25 is the presence of sterol densities within the inward-facing cavity. These sterols, which may be endogenously extracted from Sf9 cells, could potentially function as regulatory molecules for Arabidopsis AtABCG25. While sterols have been extensively studied in animals (Telbisz et al., 2014), their functions within plants remain to be further investigated.

Our study reveals the hydrophobic interactions governing ABA binding to AtABCG25 and identifies specific amino acids critical for this interaction. Notably, these key residues do not exhibit a high degree of conservation among ABA transporters in *Arabidopsis*, suggesting the existence of distinct mechanisms for ABA recognition and transport across diverse ABCG transporters. Furthermore, the structural analysis of the ATP-bound form of AtABCG25 reveals conformational changes. ATP-induced NBD closure is transmitted to the TMD, resulting in a series of structural transitions that facilitate ABA translocation. This mechanism aligns with the well-established transport cycle of ABC transporters observed across various life forms. Nevertheless, a complete understanding of the ABA release process necessitates the determination of an outward-facing ATP-bound structure. In conclusion, our study contributes valuable insights into the mechanisms governing ABA transport by AtABCG25. It provides a foundation for further exploration of sterol functions in plant transporters, the diverse substrate recognition mechanisms of ABCG transporters, and the complete transport cycle of AtABCG25-mediated ABA transport. During preparation of this manuscript, two AtABCG25 related structural studies were reported (Huang et al., 2023; Ying et al., 2023). Compared to the two published papers, our cryo-EM structures reveal conformational changes in AtABCG25 upon ABA binding compared to the apo state. Additionally, within the extracellular density, we observed intramolecular disulfide bonds in the AtABCG25 molecule. This comprehensive understanding promises to advance our knowledge of how plant tackle environmental challenges via diverse transporters-mediated hormone movement, paving the way for future crop breeding with improved resilience and stress tolerance.

## Methods

### Expression and purification of AtABCG25

The gene of *AtABCG25* (Uniprot: Q84TH5), obtained from Arabidopsis cDNA, was inserted into a modified pFastBac vector with a 3× Flag tag and a PreScission Protease cleavage site at the N-terminus. The E232Q mutation was introduced using the Fast Mutagenesis Kit (Vazyme). AtABCG25 and AtABCG25(E232Q) were expressed using the baculovirus system (Invitrogen). Protein production involved baculovirus infection of Sf9 insect cells, followed by cell harvesting, rapid freezing in liquid nitrogen, and storage at −80°C.

For protein purification, the storage cells were re-suspended in buffer A (50 mM HEPES-Na pH 7.5, 300 mM NaCl, 2 mM MgCl_2_, 10% glycerol, 1.0% LMNG, 0.2% CHS) supplemented with cocktail (MCE) and benzonase (Yeasen) and incubated at 4 °C for 2 hours. In the case of purified apo AtABCG25, CHS was removed in buffer A. After incubation, the supernatant was isolated by centrifugation at 80,000 × *g* for 1 h and incubated with Anti-DYKDDDDK G1 Affinity Resin (GenScript) at 4 °C for 30 min. The resin was washed with buffer B (50 mM HEPES-Na pH 7.0, 70 mM KCl, 2 mM MgCl_2_, 0.02% GDN) and then eluted with buffer B plus 200 μg ml^−1^ Flag peptide. The protein eluent was concentrated using a 100-kDa cut-off Centricon (Millipore) and subsequently loaded onto a Superose 6 Increase 5/150 GL column (Cytiva) equilibrated with buffer B. The fractions corresponding to the peak were combined and further concentrated to a final concentration of 6.8 mg ml^−1^ before cryo-EM sample preparation. All protein purification steps were conducted at 4 °C.

### Cryo-EM sample preparation and data collection

For cryo-EM sample preparation, 3 μl protein was applied to glow-discharged carbon grids (Quantifoil Cu R1.2/1.3 300 mesh). Subsequently, the grids were subjected to blotting for 4 s using grade 597 filter paper (Whatman) and promptly plunge-frozen in liquid ethane, which had been pre-cooled by liquid nitrogen, utilizing a Vitrobot (Thermo Scientific) maintained at 8 °C with 100% humidity. For ABA bound cryo-EM sample preparation, 0.2 mM ABA was introduced into the AtABCG25 sample, and the reaction was carried out at 25°C for 15 minutes. For E232Q mutant, protein sample was incubated with 0.2 mM ABA, 5 mM ATP and MgCl_2_ at 25°C for 15 minutes prior to the grid freezing step.

For data collection of CHS-bound AtABCG25 and ABA-bound AtABCG25, the grids were loaded into a Titan Krios (FEI) electron microscope operating at 300 kV and equipped with the energy filter and K3 direct electron detector (Gatan). Movies stacks recorded using SerialEM software (Mastronarde, 2005) in the super-resolution mode, with a calibrated pixel size of 0.55 Å at a nominal magnification of ×105,000. Each stack was acquired with a total electron dose of 50 e^−^ Å^−2^ for 32 frames and a defocus range from −0.8 to −1.8 μm. For data collection of apo AtABCG25 and ATP-bound AtABCG25(E232Q), movies stacks recorded using EPU software in the super-resolution mode, with a calibrated pixel size of 0.5475 Å at a nominal magnification of ×81,000.

### Data processing

The data processing workflow is illustrated in Supplemental Figure 2. Motion correction and dose weighting were performed using the EMshark implementation of MotionCor2 (Zheng et al., 2017) and the stacks were binned twofold, resulting in a pixel size of 1.1 Å or 1.095 Å. The dose-weighted and motion-corrected micrographs were then imported into cryoSPARC (Punjani et al., 2017) to determine their defocus values and CTF parameters via Patch CTF Estimation. Particles pick by blob picker or template picker, and then extract for 2D classification. Junk particles were removed by several rounds of 2D classification. The initial 3D model was generated using ab initio reconstruction. Junk particles were also removed by ab initio reconstruction and heterogeneous refinement. The best classes were used for non-uniform refinement in order to enhance the resolution. Non-uniform refinement was carried out with both C1 and C2 symmetries. The overall resolution of each map was determined by the gold-standard FSC= 0.143 criterion. Local resolution estimation for each map was generated using cryoSPARC.

### Model building and structure refinement

The initial monomer model of AtABCG25 was generated using AlphaFold2 (Tunyasuvunakool et al., 2021), and dock into the cryo-EM maps using UCFC Chimera (Pettersen et al., 2004). Subsequent to this, manual model building, and adjustment were performed using Coot (Emsley and Cowtan, 2004). ABA, ATP, and magnesium ions were obtained from the ligand library within Coot. The refinement of the structures was carried out through real-space refinement using PHENIX (Adams et al., 2010). All structure figures were prepared in PyMOL (https://pymol.org/2/), and ChimeraX (Pettersen et al., 2021).

### ATPase activity assay

The ATPase activity was assessed with slight modifications to a previously published protocol (Mu et al., 2023). ATPase activity was measured using a commercially available kit that quantified the release of inorganic phosphate (Pi) from ATP following the kit’s instructions (Nanjing Jiancheng Bioengineering Institute, China). In brief, enzymatic reactions were carried out for 10 minutes at 37°C in a final volume of 68 μL, containing various concentrations of ATP. The purified WT or mutant AtABCG25 (0.95 μg) in a buffer containing of 50 mM HEPES-Na pH 7.0, 70 mM KCl, 2 mM MgCl_2_, 0.02% GDN were uesd. The reactions were then mixed with 42 μL of matrix reagent for an additional 10 minutes at 37°C and terminated by adding 10 μL of reagent buffer. After centrifugation at 3000×g for 10 minutes, the supernatant was collected for Pi content measurement. A total of 30 μL of the supernatant was transferred to a 96-well plate, mixed with 100 μL of chromogenic agent for 2 minutes at room temperature, and then followed by the addition of 100 μL of Reagent 6 for 5 minutes. Absorbance was measured at 636 nm using the microplate reader (Tecan, Switzerland). Nonlinear regression to the Michaelis–Menten equation and statistical analysis was performed using GraphPad Prism version 8.4.0.

### ABA sensor-based cellular transport assays

Tobacco mesophyll protoplasts were extracted and transfected as previously described (Zhou et al., 2021) with a slight difference. Briefly, tobacco protoplasts were either transfected with the cytosolic ABA sensor ABAleon_Tao3 (Tao3_cyto_) alone or co-transfected with ABCG25 or its mutants (I445A, V449A, L556A). After transfection, cells were incubated at 25℃ for 16-24 h and treated with ABA (1 μM) or incubation buffer (mock) for 3 h before imaging. Fluorescence lifetime imaging microscopy (FLIM) recoding of the ABA sensor was performed data were analyzed as previous (Zhou et al., 2021). For each group, at least 10 cells were imaged and recorded by FLIM.

## Acknowledgements

We would like to thank the staffs at SUSTech Cryo-EM Center for help in data collection. This work was supported by National Natural Science Foundation of China (32271251 to K.Y., 32070292 to J.L), Guangdong Innovative and Entrepreneurial Research Team Program (2021ZT09Y104, 2021QN02Y429 to K.Y.), Shenzhen Science and Technology Program (No. JCYJ20220530115214033 and No. KQTD20210811090115021 to K.Y., JCYJ20170817104523456 and KQTD20190929173906742 to J.L), and Scientific research funding for postdoctoral researchers staying at Shenzhen (K20227507 to Y.Z). K.Y. is an investigator of SUSTech Institute for Biological Electron Microscopy.

## Conflict of interest statement

None declared.

## Author contributions

J.X., Y.Z. and Y.Q. performed the experiments. J.X., Y.Z., Y.Q., H.G., J.L., Y.S. and K.Y. contributed to data collection, structural determination, analysis and discussion. J.X., J.L., Y.S. and K.Y. conceived the study and wrote the paper.

## Data availability

The 3D cryo-EM density maps of ABCG25 have been deposited in the Electron Microscopy Data Bank (EMDB) under the accession numbers EMD-37418, EMD-37426, EMD-37397 and EMD-37455, respectively. Coordinates for structure models have been deposited in the Protein Data Bank (PDB) under the accession codes 8WAM, 8WBA, 8WBX and 8WD6, respectively.

---

**Supplemental Fig.1.**
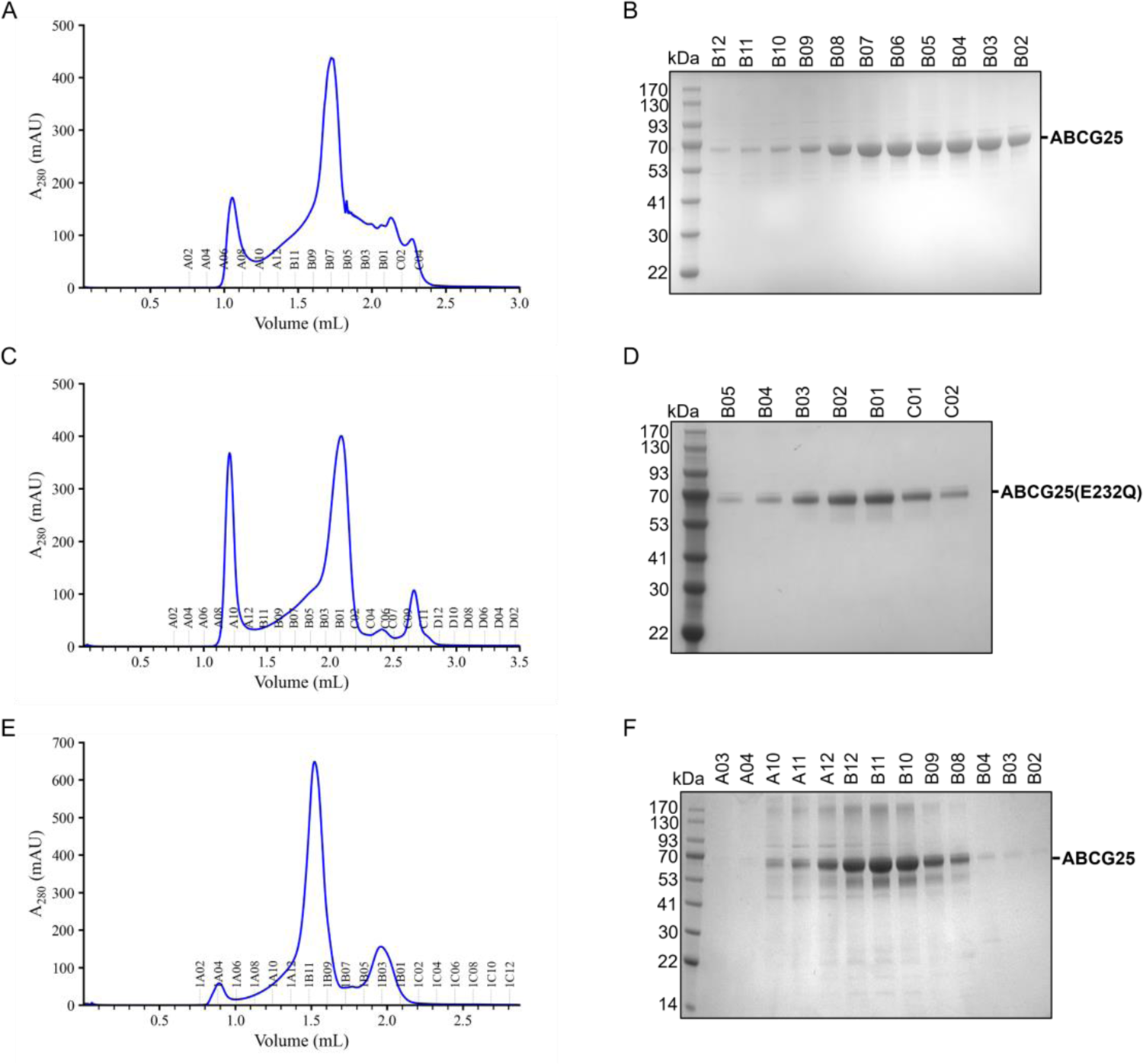
Purification of AtABCG25. (A) Gel filtration of AtABCG25, extracted with LMNG+CHS, was conducted using a Superose 6 Increase 5/150 GL column in a buffer containing 0.02% GDN. (B) The SDS-PAGE gel result of the protein corresponding to the peak position in Figure A. (C) Gel filtration of AtABCG25(E232Q), extracted with LMNG+CHS, was conducted using a Superose 6 Increase 5/150 GL column in a buffer containing 0.02% GDN. (D) The SDS-PAGE gel result of the protein corresponding to the peak position in Figure C. (E) Gel filtration of AtABCG25, extracted with LMNG, was conducted using a Superose 6 Increase 5/150 GL column in a buffer containing 0.02% GDN. (F) The SDS-PAGE gel result of the protein corresponding to the peak position in Figure A.

**Supplemental Fig. 2.**
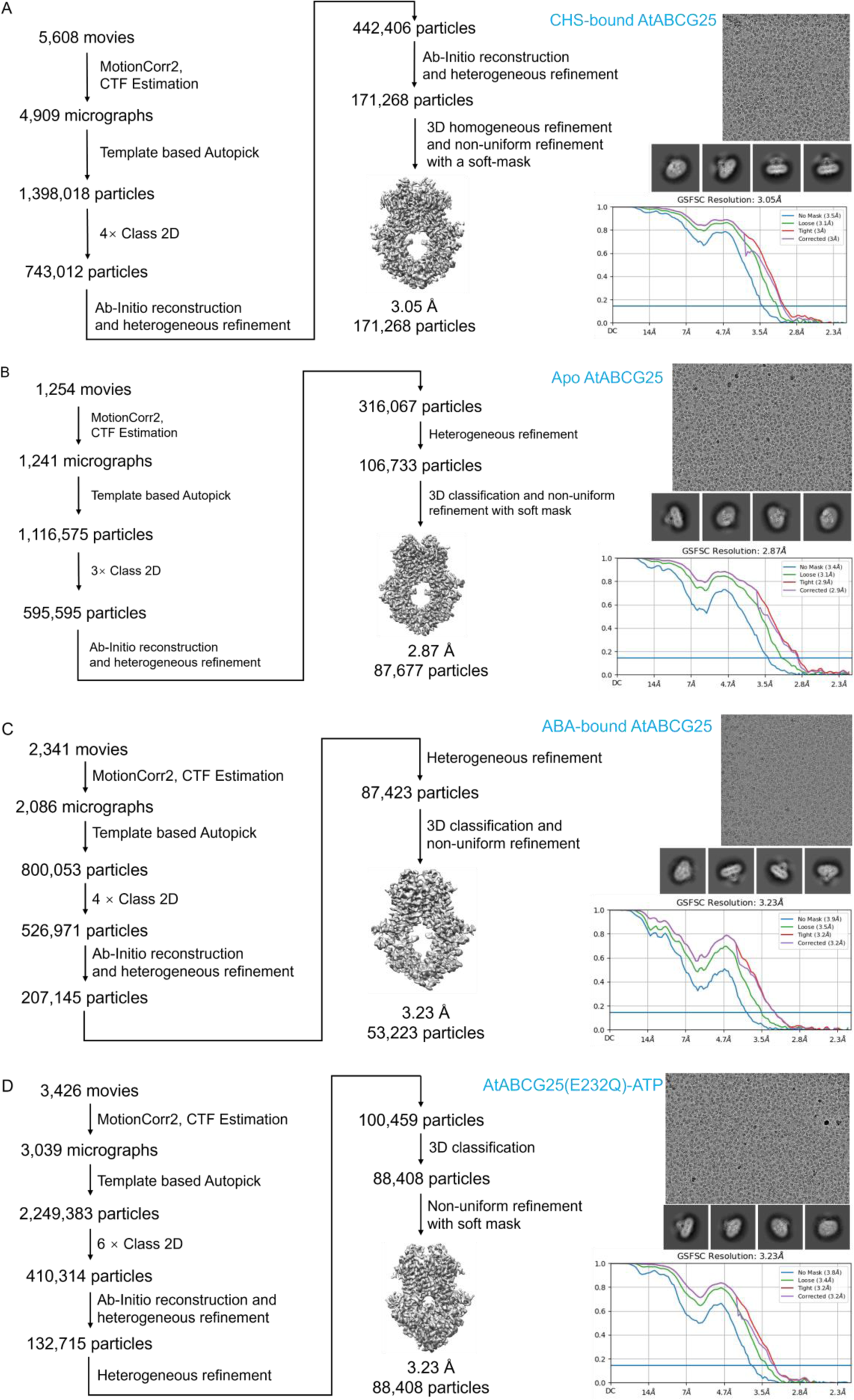
Cryo-EM data processing of AtABCG25. (A) Flowchart of the data processing, representative micrograph, 2D class averages and FSC curve for CHS bound to AtABCG25. (B) Flowchart of the data processing, representative micrograph, 2D class averages and FSC curve for apo AtABCG25. (C) Flowchart of the data processing, representative micrograph, 2D class averages and FSC curve for ABA bound to AtABCG25. (D) Flowchart of the data processing, representative micrograph, 2D class averages and FSC curve for ATP bound to AtABCG25(E232Q).

**Supplemental Fig. 3.**
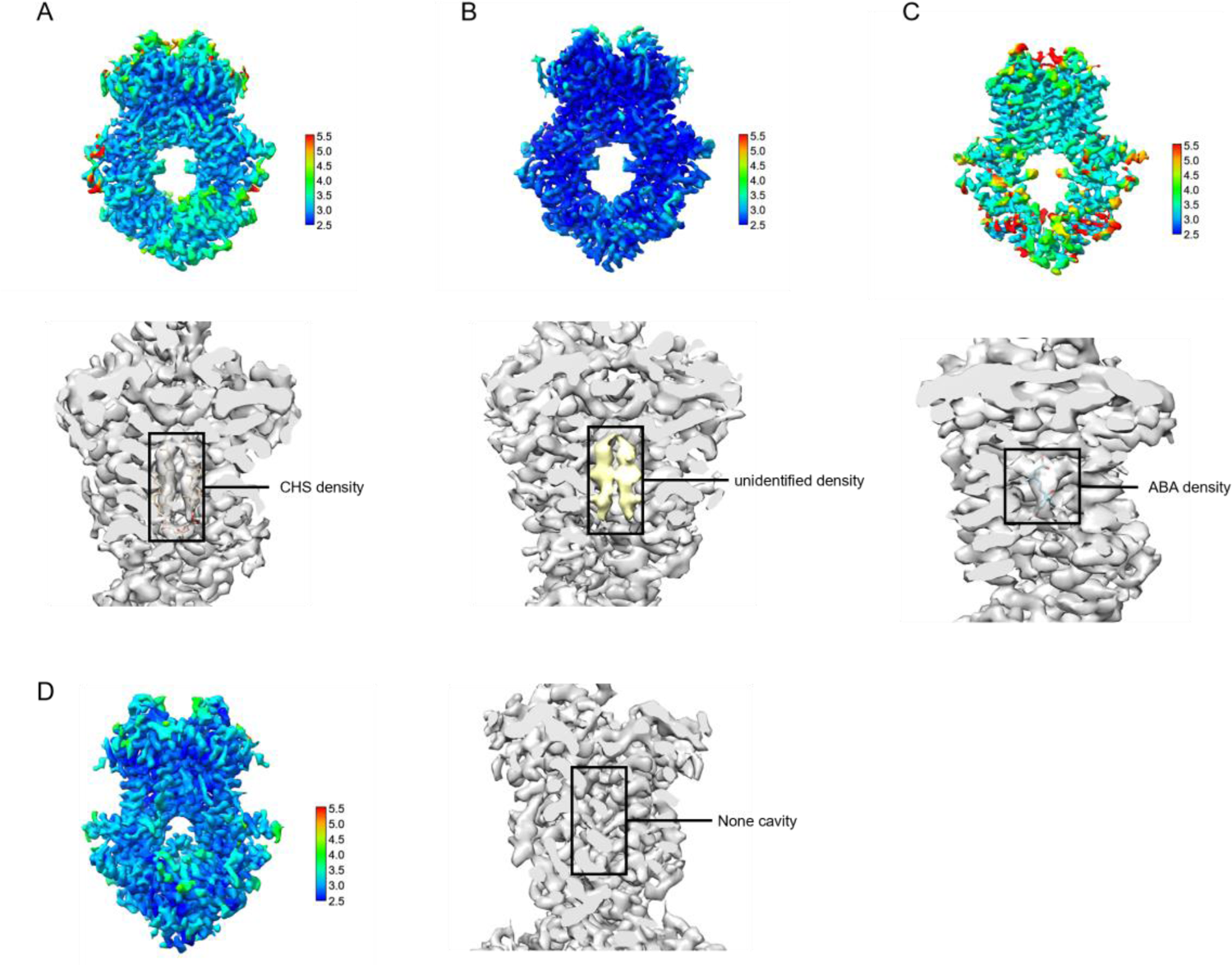
Local resolution estimation of AtABCG25. (A) Local resolution estimation of CHS bound to AtABCG25 and CHS density. (B) Local resolution estimation of apo AtABCG25 and unidentified density. (C) Local resolution estimation of ABA bound to AtABCG25 and ABA density. (D) Local resolution estimation of ATP bound to AtABCG25(E232Q).

**Supplemental Fig. 4.**
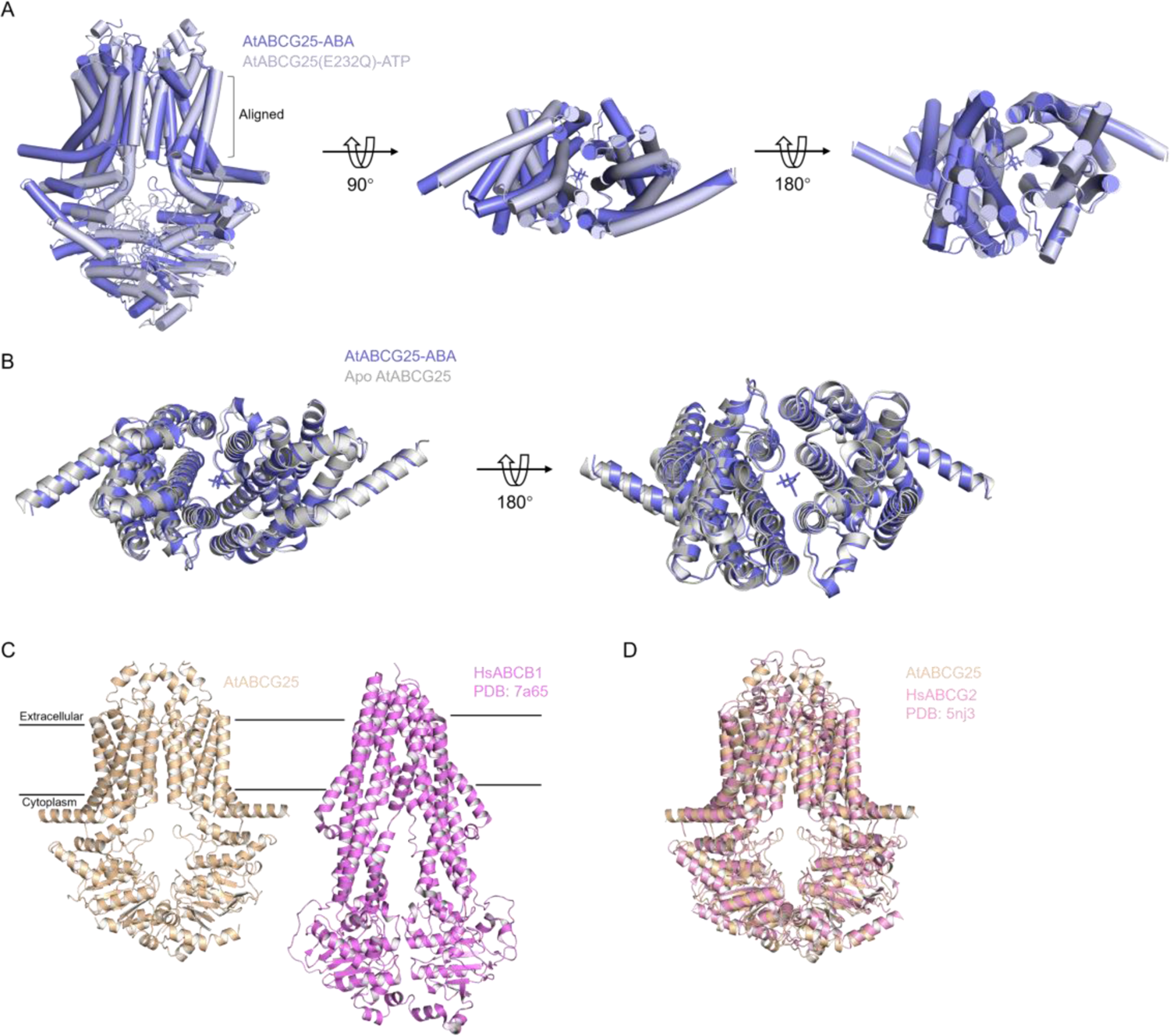
Structural alignment of AtABCG25. (A) Structural alignment between ATP-bound AtABCG25(E232Q) and ABA-bound AtABCG25 across the TMD. ATP-bound AtABCG25(E232Q) is depicted in bluewhite, while ABA-bound AtABCG25 is shown in slate. Residues not used are hidden. (B) Other views of structural alignment between apo AtABCG25 and ABA-bound AtABCG25 across the TMD. Apo AtABCG25 is depicted in gray, while ABA-bound AtABCG25 is shown in slate. Residues not used are hidden. (C) Overall structure of AtABCG25 (colored wheat) and HsABCB1 (colored violet). (D) Structural alignment between AtABCG25 (colored wheat) and HsABCG2 (colored pink).

**Supplemental Fig. 5:**
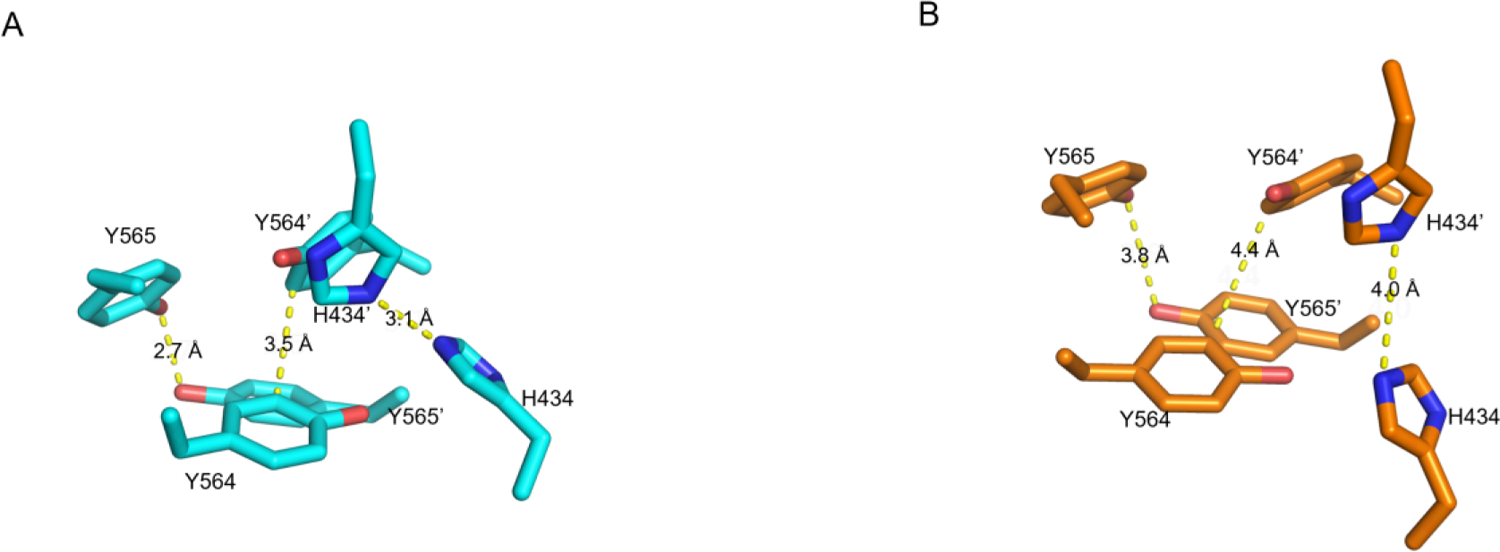
Changes in the distance of Y564, Y565, and H434. (A) The distances between Y564 and Y564’, Y565 and Y565’, and H434 and H434’ in the closed state (inward). (B) The distances between Y564 and Y564’, Y565 and Y565’, and H434 and H434’ in the open state (outward).

**Supplemental Table 1:** Summary of cryo-EM data collection, processing, model refinement and validation.

| **Data collection** | Apo<br>AtABCG25 | AtABCG25-CHS | AtABCG25-ABA | AtABCG25EQ-ATP |
| --- | --- | --- | --- | --- |
| EM equipment | Titan Krios | Titan Krios | Titan Krios | Titan Krios |
| Voltage (kV) | 300 | 300 | 300 | 300 |
| Detector | Gatan K3<br>Summit | Gatan K3<br>Summit | Gatan K3<br>Summit | Gatan K3 Summit |
| Pixel size (Å) | 1.095 | 1.1 | 1.1 | 1.095 |
| Total electron dose (e<sup>-</sup><br>Å<sup>-2</sup>) | 50 | 50 | 50 | 50 |
| Defocus range (μm) | -0.8 ~ -1.8 | -1.0 ~ -2.0 | -0.8 ~ -1.8 | -0.8 ~ -1.8 |
| **3D Reconstruction** |  |  |  |  |
| Software | CryoSPARC | CryoSPARC | CryoSPARC | CryoSPARC |
| Number of micrographs | 1254 | 5608 | 2341 | 3426 |
| Final particles | 87677 | 171268 | 53223 | 88408 |
| Symmetry | C2 | C2 | C2 | C2 |
| Map resolution (Å) | 2.87 | 3.05 | 3.23 | 3.23 |
| FSC Threshold | 0.143 | 0.143 | 0.143 | 0.143 |
| Map sharpening B-factor (Å<sup>2</sup>) | -96.1 | -124.1 | -107.7 | -120.9 |
| **Refinement and**<br>**validation** |  |  |  |  |
| Protein residues | 1025 | 1106 | 918 | 1102 |
| Ligand | 0 | 2 | 1 | 4 |
| R.M.S. deviations |  |  |  |  |
| Bond lengths (Å) | 0.002 | 0.002 | 0.003 | 0.002 |
| Bond angles (°) | 0.563 | 0.537 | 0.631 | 0.449 |
| MolProbity score | 1.43 | 1.50 | 1.51 | 1.31 |
| Clash score | 5.38 | 9.38 | 9.41 | 5.64 |
| Rotamer outliers | 0.46 | 0.32 | 0.38 | 0.11 |
| Ramachandran plot (%) |  |  |  |  |
| Preferred | 97.31 | 98.43 | 98.26 | 98.34 |
| Allowed | 2.69 | 1.57 | 1.74 | 1.66 |
| Outliers | 0 | 0 | 0 | 0 |

## Notes

### Competing Interest Statement

The authors have declared no competing interest.

